# Spectral clustering in regression-based biological networks

**DOI:** 10.1101/651950

**Authors:** Sheila M. Gaynor, Xihong Lin, John Quackenbush

## Abstract

Biological networks often have complex structure consisting of meaningful clusters of nodes that are integral to understanding biological function. Community detection algorithms to identify the clustering, or community structure, of a network have been well established. These algorithms assume that data used in network construction is observed without error. However, oftentimes intermediary analyses such as regression are performed before constructing biological networks and the associated error is not propagated in community detection. In expression quantitative trait loci (eQTL) networks, one must first map eQTLs via linear regression in order to specify the matrix representation of the network. We study the effects of using estimates from regression models when applying the spectral clustering approach to community detection. We demonstrate the impacts on the affinity matrix and consider adjusted estimates of the affinity matrix for use in spectral clustering. We further provide a recommendation for selection of the tuning parameter in spectral clustering. We evaluate the proposed adjusted method for performing spectral clustering to detect gene clusters in eQTL data from the GTEx project and to assess the stability of communities in biological data.

## 1 Introduction

Networks are increasingly used to represent the complex patterns of relationships between biological features. Interactions between genetic variants, genes, proteins, and other biological data sources can be captured in a single, flexible framework as networks [1, 2]. Expression quantitative trait loci (eQTL) mapping yields associations between single nucleotide polymorphisms (SNPs) and gene expression levels. Representing eQTLs as bipartite networks provides insight into regulatory function and interactions that drive phenotypes [3, 4]. The structure and metrics of eQTL networks have elucidated functionality and have been demonstrated to be highly modular. In multiple settings, genome-wide significant SNPs have been shown to be central to communities rather than network-wide hubs. These communities are defined by groups of SNPs and genes that have been annotated with common function and thus likely act in concert to regulate function. Polygenic networks can also be constructed from the SNP-gene associations to further inform biological function from network structure with a specific focus on gene-gene interactions.

The construction of eQTL networks generally proceeds by (a) defining an adjacency matrix from the results of a linear regression of gene expression on additive SNP genotype, and then (b) performing community detection on the adjacency matrix to identify groupings of nodes. Many community detection approaches have been proposed towards identifying highly connected, clustered nodes in networks [5]. For instance, one may use a modularity maximization approach that proceeds to group nodes such that there are more edges within the community than would be expected by chance. [4, 6]. This approach is widely employed to ensure that nodes within a community are densely connected whereas those in different communities are sparsely connected. Modularity maximization, however, often can fail to detect small communities. Hierarchical clustering is also used for community detection to group similar nodes based upon a defined similarity matrix, resulting in a tree-like structure that decomposes the data into hierarchies of clusters [7, 8]. This approach is highly sensitive to the chosen similarity measure and lacks robustness when clustering noisy biological data [8, 9]. Lastly, one may use spectral clustering to cluster nodes based on their affinity to one another [10–15].

Spectral clustering is an algorithm used for community detection that has been widely applied, including for biological data such as gene expression and protein levels [16–20]. Spectral clustering is well suited to biological data as it maps the network to a low-dimensional space and then detects communities. In particular, the approach operates on an affinity matrix representing the likeness of data points to one another. The top eigenvalues of a Laplacian matrix of the affinity matrix are then used in standard clustering procedure, such as k-means clustering [21]. This, in effect, allows the method to identify communities in the network in lower dimension. Spectral clustering is efficient, highly flexible, and does not rely on standard distance metrics.

Community detection approaches, including spectral clustering, proceed with the assumption that the network’s adjacency matrix is the fixed truth [22]. In certain circumstances in network sciences, such as computer and social networks, the adjacency matrix can be exactly observed [23]. However, in many biological settings the adjacency matrix is estimated from preliminary analyses. For instance, in eQTL networks the adjacency matrix is derived as a function of the associations between SNPs and genes as estimated in linear regression. Community detection approaches thereby neglect the fact that data inputs have been previously estimated, which is notable as error results from statistical estimation. This error is thus not propagated in the proceeding network analysis of community detection. Previous work by Huang et al [2009] demonstrated the effect of perturbations on spectral clustering performance, showing the influence of network nodes measured with error [24]. They evaluated the propagation of error in a stepwise manner, from data error to the mis-clustering rate, and formalized the loss of clustering performance.

In this paper we characterize the impact of performing community detection when the adjacency matrix was initially estimated in a regression analysis rather than directly observed. We first describe the spectral clustering algorithm as applies to gene network constructed from an eQTL analysis. We then characterize the impact of error in the network’s adjacency matrix with respect to the affinity matrix and Laplacian matrix. We consider a simple adjustment to account for the bias in the affinity matrix and apply it to data from the Genotype-Tissue Expression (GTEx) consortium. We assess the impact on detection of Gene Ontology sets and reproducibility of clusters. The paper concludes with discussion of the method, applications and future work.

## 2 Results

We evaluated the community structure of eQTL networks built from data from the NHGRI Genotype-Tissue Expression (GTEx) project [25]. The GTEx project is a resource of genotype and expression data from multiple human tissues from hundreds of human donors. We used the version 6.0 imputed genotyping and RNA-seq data from dbGaP. The RNA-Seq data were preprocessed using the YARN package in R and normalized using the qsmooth package in R [26, 27]. We included tissues that had over 200 observations and are annotated in the Roadmap Epigenomics Project core 15-states model, yielding 11 tissues. This selection was made to identify and include only SNPs that are in active chromatin regions in order to reduce the SNP space and increase the likelihood of signal in the adjacency matrix.

We sought to identify clusters of genes in the GTEx data. Fagny et al [2017] observed that different communities in eQTL networks are enriched for different Gene Ontology (GO) biological process gene sets. We thus considered clustering data comprised exclusively of two distinct sets of GO biological processes in each tissue. In particular, we clustered all genes available from pairs of GO biological process sets that (a) were identified to be enriched in distinct communities in the Fagny et al [2017] and (B) were disjoint sets not sharing any genes in common. This selection was made in order to ensure identifiability and prevent GO sets from clustering together. The data were further limited to sets with at least ten and less than 500 genes in order to ensure they had specific functionality. The cluster setting is defined for all the tissues in Table 1, with the sample size available in each tissue and GO set pairings available. We then sought to identify the two clusters of GO biological process gene sets using spectral clustering.

**Table 1:** GO biological process gene set pairings (i.e. number of clustering settings) considered in each GTEx tissue.

| Tissue | Sample size | Pairs of gene sets |
| --- | --- | --- |
| Adipose - subcutaneous | 313 | 2801 |
| Artery - aorta | 216 | 1918 |
| Breast - mammary | 185 | 266 |
| Cells - transformed fibroblasts | 291 | 3260 |
| Esophagus - mucosa | 276 | 2955 |
| Heart - left ventricle | 178 | 18972 |
| Lung | 290 | 6 |
| Muscle - skeletal | 378 | 6109 |
| Pancreas | 168 | 865 |
| Skin | 370 | 24794 |
| Whole blood | 365 | 696 |

Community detection was performed to detect gene communities for all GO set pairings in each tissue in Table 1 in two ways: (1) using the standard spectral clustering algorithm of Ng et al [2002] and (2) using the proposed spectral clustering with adjustment in the affinity matrix to account for the biased induced by intermediate regression analyses as described in the Methods. In both cases, the number of clusters was fixed to be equal to two and the scaling parameter was selected as described in the Methods. The concordance of the communities of genes detected by each spectral clustering method and the true GO biological process communities was assessed by the Kappa statistic, a measure of agreement between group assignments. Specifically, having obtained the group assignments a 2 × 2 concordance table was constructed with rows and columns ordered to maximize the trace of the table in order to account for the arbitrary group labels. The weighted kappa statistic was then computed based upon these assignments.

Figure 1 demonstrates the difference between the Kappa statistic calculated from the spectral clustering approach adjusted for error and the Kappa statistic calculated from the standard spectral clustering algorithm. In each of the tissues the difference in Kappa statistics is centered around zero, demonstrating that the algorithms detect clusters concordant with the GO sets similarly in these settings. We performed a one-sided test in order to evaluate if the average difference in Kappa statistics is greater than zero in each of the tissues, which would suggest that the approach adjusting for error yields clusters more concordant with the GO sets. For every tisue, we do not obtain a statistically significant result. Thus on average we do not observe a difference in the ability of the two spectral clustering approaches considered to detect clusters defined by the two Gene Ontology sets.

**Figure 1:**
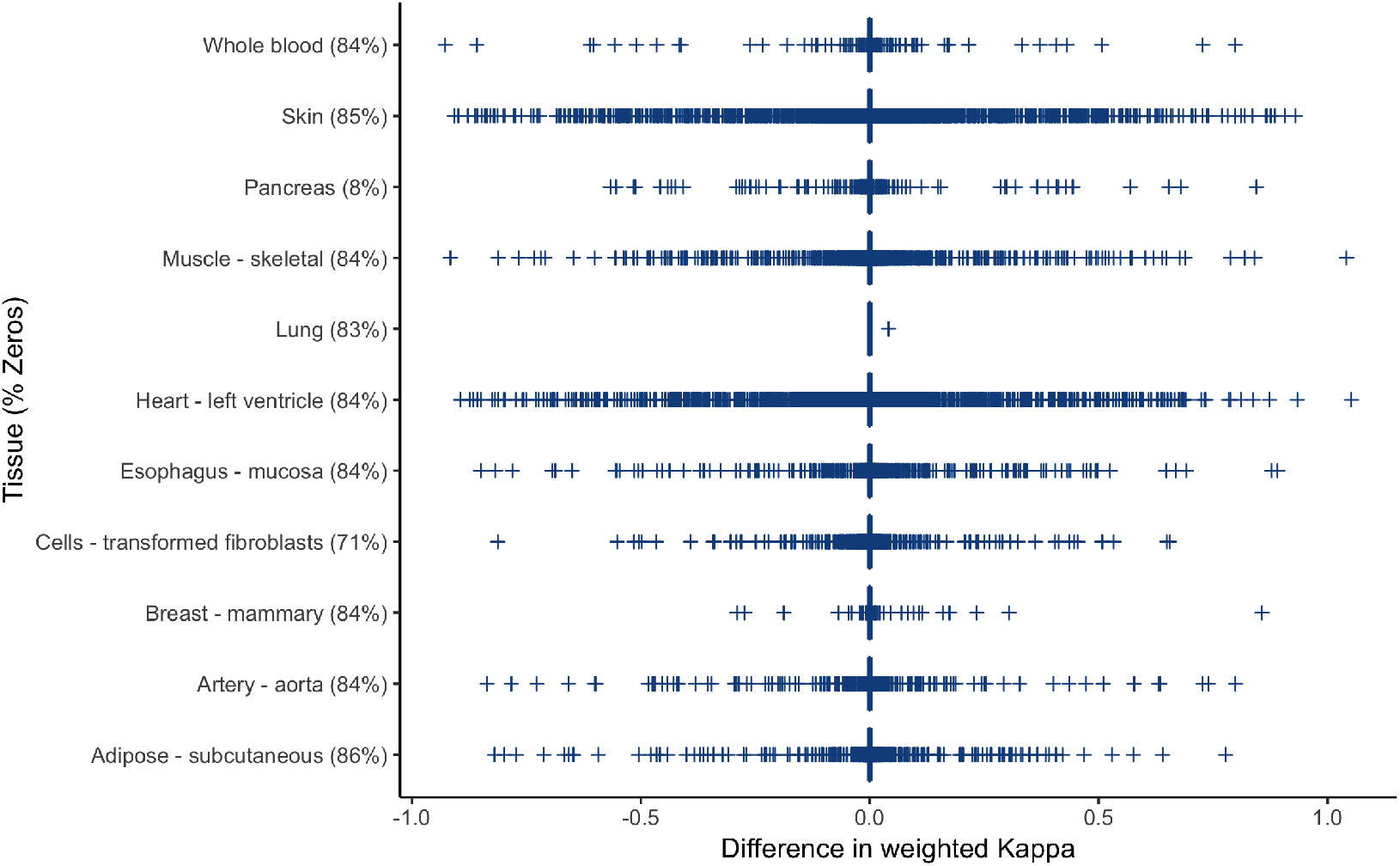
Difference between the weighted Kappa statistic for the concordance of the communities detected via standard spectral clustering and spectral clustering with error adjustment.

We next assessed the reproducibility of community detection by each of the spectral clustering approaches. We restricted our analysis to the setting where clusters identified via spectral clustering were highly enriched for GO sets. Specifically, we selected the pair of GO sets in each tissue with the highest concordance to the true GO sets. The selected GO sets for each tissues are described in Table 2. For each of these settings, we randomly split the sample such that half of the observations are assigned to be a training set while the remaining half are considered the test set. In both the training and tests sets, we perform eQTL mapping and community detection to identify groupings of genes. Community detection was performed again using both spectral clustering methods. The Kappa statistic computed to assess the concordance between the gene communities in the training and test sets in each method. The procedure of data splitting, obtaining group or community assignments for genes, and calculating the weighted kappa statistic was repeated for 500 random data splits in order to account for variation in the sample splitting.

**Table 2:** GO sets analyzed via sample splitting for concordance between clusters identified from standard and adjusted spectral clustering.

| Tissue | Gene Ontology sets | $n_1$ | $n_2$ |
| --- | --- | --- | --- |
| Adipose - subcutaneous | response to interferon-gamma & DNA conformation change | 137 | 210 |
| Artery - aorta | cytokine activity & lysosome membrane | 157 | 263 |
| Breast - mammary | homophilic cell adhesion via plasma membrane adhesion molecules & coated vesicle membrane | 148 | 128 |
| Cells - transformed fibroblasts | translational initiation & intermediate filament | 175 | 95 |
| Esophagus - mucosa | GTP binding & carbohydrate binding | 336 | 228 |
| Heart - left ventricle | positive regulation of T cell activation & proton transmembrane transport | 178 | 92 |
| Lung | homophilic cell adhesion via plasma membrane adhesion molecules & intermediate filament | 146 | 101 |
| Muscle - skeletal | integral component of organelle membrane & contractile fiber | 138 | 209 |
| Pancreas | chromatin assembly & peptide binding | 97 | 196 |
| Skin | cellular modified amino acid metabolic process & negative regulation of cell cycle process | 141 | 206 |
| Whole blood | T cell receptor signaling pathway & chromatin assembly or disassembly | 152 | 124 |

The results of this analysis are given in Table 3. We observe that there is no significant improvement in concordance between splits by adjusting for the bias in the affinity matrix when seeking to identify consistent clusters. We observe similar concordance between sample splits regardless of the affinity matrix employed in spectral clustering. We note the decreased sample sizes in the splits could lead to greater error in the initial eQTL mapping, as the sample sizes ranged from 108 to 189 in each set considered for clustering when halved.

**Table 3:** Concordance of clusters identified in settings of pairs of GO biological processes between split samples.

| Tissue | Standard Clustering | Adjusted Clustering |
| --- | --- | --- |
| Adipose - subcutaneous | 0.99 (0.09) | 1.00 (0.04) |
| Artery - aorta | 1.00 (0) | 1.00 (0) |
| Breast - mammary | 0.98 (0.15) | 0.98 (0.13) |
| Cells - transformed fibroblasts | 0.76 (0.23) | 0.75 (0.23) |
| Esophagus - mucosa | 1.00 (0) | 1.00 (0) |
| Heart - left ventricle | 0.77 (0.23) | 0.75 (0.23) |
| Lung | 1.00 (0) | 1.00 (0) |
| Muscle - skeletal | 1.00 (0.04) | 0.99 (0.12) |
| Pancreas | 0.99 (0.06) | 0.98 (0.10) |
| Skin | 1.00 (0.04) | 0.99 (0.10) |
| Whole blood | 0.99 (0.11) | 0.98 (0.13) |

## 3 Discussion

Community detection methods to identify groups of nodes that are more connected or related to each other have been highly explored in many network settings. They are particularly useful in biological networks, where clusters of nodes often indicate shared biological function or important pathways. Spectral clustering is an approach to community detection that straightforwardly allows one to detect community structure in a lower-dimensional space than the data. In this work, we consider the specific application of spectral clustering to identify clusters of genes in eQTL networks. In such a setting, the adjacency matrix defining the network has been estimated through prelimary regression analyses and therefore have associated error. We characterize the impact of error when performing the spectral clustering method to affinity matrices of eQTL networks. We flexibly allow the errors for each observation to have their own variance specification, moving beyond the simple setting of normal noise perturbing the complete data set.

Given our formulation of the error on the data used in spectral clustering, we proposed a simple adjustment for the error. We consider the practice of performing spectral clustering on an unbiased affinity matrix, which allows for one to account for the additional estimation step that occurs when a network is built upon intermediate regression results rather than observed or true values. We apply this approach to a setting where we anticipated finding two distinct gene clusters in eQTL data. The success in identifying the two gene clusters within all settings varied by tissue and on average we did not see that including such an error adjustment improved our ability to detect such structure. Further, we did not observe a difference in the ability to consistently detect communities. This suggests a limited impact of the estimation of eQTLs in these networks; it would be of value to explore settings with intermediary regression analyses accross different effect sizes, error, and sample sizes.

We additionally introduce a simplification of the local scaling approach for selection of the tuning parameter in the spectral clustering approach, as described in the Methods. The implementation of spectral clustering by Ng, Jordan, and Weiss [2002] and similar approaches rely upon the user to input the parameter, and oftentimes this can highly influence the clustering results. The complete self-tuning local scale approach of Zelnik-Manor and Perona [2005] provides a clear framework for approaching the selection of the parameter using the data. The current implementation, however, increases the computational burden by requiring a different parameter estimate for each data point. We observe in our data setting that estimated local scaling parameters do not follow a fat-tailed distribution, and propose to take a summary of the local parameters for application across all data points. This general framework would also apply if one chose to use a different cost function for the distance metric. This contribution allows for more efficient, data-driven spectral clustering.

Our evaluation of the impact of error is naturally extended to other biological networks that are constructed on results from intermediary analyses. For instance, one could use methylation data in place of gene expression in order to obtain mQTL networks and identify gene communities with shared function. In such QTL settings, we obtain a regression-based network with error of the same form as described in the Methods, given that one is using the coefficient estimates in the adjacency matrix. Further, This is more broadly applicable outside of biological networks to settings where the observed network has error that can be estimated and assumed to follow a normal distribution. This lends itself towards future work in assessing whether more sophisticated clustering approaches are better suited to incorporating measures of error for such settings and characterizing the impact of error and ability to easily adjust in other biological network settings.

## 4 Materials and methods

### 4.1 Setting and algorithm

We first introduce the steps of the spectral clustering algorithm as it would standardly proceed to identify community structure. We present the setting where the network is based upon regression coefficients in order to apply to the setting of eQTL networks, where we are interested in communities of genes. The spectral clustering approach can be applied more broadly to any setting in which one wishes to detect communities based upon a set of features defining a matrix representing the network, termed the adjacency matrix.

In the eQTL setting, we estimate a regression coefficient to represent the association between SNP genotypes and gene expression. Suppose that one fits the following regression model between some exposure *S*_*i*_ and outcome *G*_*j*_ for a set of independent observations, *G*_*j*_ = **X**^**T**^***α*** + *β*_*ij*_ *S*_*i*_. In the eQTL setting, we assume a *z* × *n* matrix of SNP genotypes **S** and *z* × *m* matrix of gene expression **G**. Each of these matrices have *z* rows of observations columns containing the *n* SNPs and *m* genes, respectively. We adjust for some set of covariates **X**, such as principle components for population structure, sex, and age. Thus our model represents an eQTL regression of a particular SNP *i* on a locus’s gene expression *j*. The associations from the regression models can then be used to define the adjacency matrix **Q**, an *m* × *n* matrix of regression coefficients *β* from the regressions of all *n* SNPs on all *m* genes, shown below.

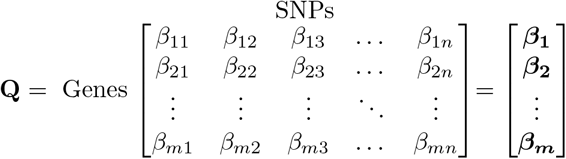

Community detection is applied to this adjacency matrix **Q** via spectral clustering. We consider the spectral clustering algorithm as given in Ng et al [2002] and described as follows. First, define the affinity matrix **A**, a representation of the similarity between different nodes in **Q**. We define **A** by the Gaussian kernel similarity function, such that *A*_*ij*_ = *exp*(−‖***β***_*i*_ − ***β***_*j*_‖^2^/*σ*^2^), *A*_*ii*_ = 0 where ***β***_*i*_ is a row of **Q**. We notes that there is a scaling parameter *σ*^2^ in this algorithm which must be specified. We return to the selection of this parameter in Section 4.4. Thus, **A** is defined as

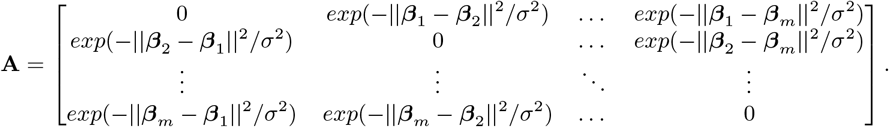

Given the affinity matrix A, we define the diagonal matrix **D** by setting *D*_*ij*_ = 0, *D*_*ii*_ = Σ_*j*_ *A*_*ij*_, i.e. define the *i*^*th*^ diagonal element to be the *i*^*th*^ row sum. This is given as

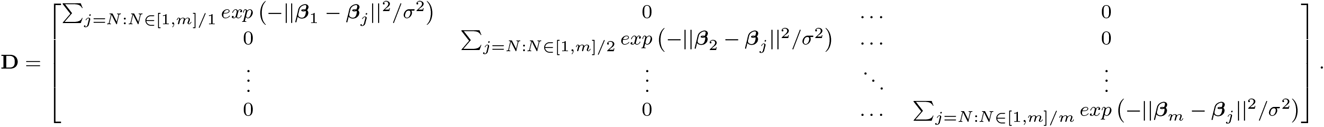

We define the Laplacian matrix **L** as **L** = **D**^−1/2^**AD**^−1/2^, which is specified as the normalized Laplacian. We then find the *k* eigenvectors of **L** with the largest absolute value, where *k* is selected based upon the eigenvalue heuristic. Define the matrix **X** to the be corresponding eigenvectors, **X** = [**x**_1_, **x**_2_,…**x**_*k*_]. The matrix **Y** is defined by normalizing the matrix **X** such that all rows have unit length, 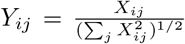. The algorithm then proceeds to cluster the matrix **Y** using k-means clustering to minimize the within cluster sum of squares. K-means clustering iterates through the following steps: (1) randomly assign k data points from **Y** as centroids, (2) placing remaining data points to closest centroids based on distance metric, (3) estimating the new cluster centroids and (4) repeating the assignment and calculation steps until a stopping criterion is met [28].

### 4.2 Error in input data

In the algorithm as defined, we proceed as though we have observed the true values of **A**. Thus for each element of **A**, we assume the following form of *A*_*ij*_ for any *i* = 1,…, *n* and *j* = 1, ‥, *m*.

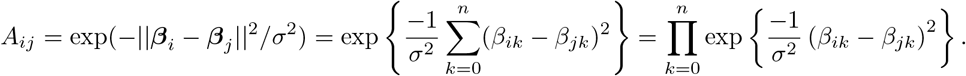

We aim to characterize the the impact of performing spectral clustering on data observed with error. We make an assumption on the form of the error in the observed 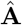. We assume that all *ε*_*ik*_ and *ε*_*jk*_ are independent and distributed as Gaussian 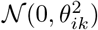 and 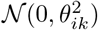. Huang et al [2009] considered the impact of perturbations on spectral clustering specifically in the specific scenario where all *ε*_*ik*_ and *ε*_*jk*_ are independent and with mean zero and bounded variance *σ*^2^. They make the distributional assumption that all errors are Gaussian when considering the effect of perturbations on the affinity matrix [13]. This formulation requires that all errors are distributed with equal variance; however, this oftentimes does not hold true. Particularly in the setting of a network derived from a large set of regression models, we wish to allow the variance to differ across the regression coefficients informing the affinity matrix.

In our setting we therefore allow the error terms to have different variance estimates across all elements of the network. For instance, in the eQTL setting we estimate the regression coefficients from the genotype and expression data to define the affinity matrix **A**. Thus for elements of **Q** (the adjacency matrix of the eQTL setting) we observe 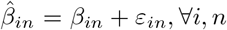. This diverges from the standard spectral clustering approach where we assume fixed values in **Q**, as we are utilizing estimates of 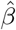 rather than true *β*. Thus rather than observing the true *A*_*ij*_, we are able to estimate 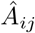. The formulation of elements of 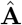 assuming they are derived from such regression models is given as

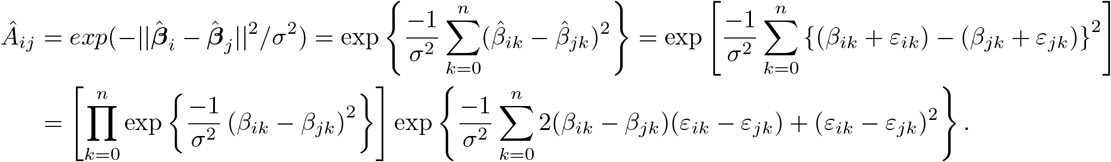

We observe from the second term that the elements of the affinity matrix are biased in this setting. We proceed to define the moments of the observed affinity matrix, maintaining our defined notation while noting that the results are generalizable to any setting with such an error structure regardless of intermediary analysis. We first consider the impact of perturbations on the affinity matrix directly in the same approach as Huang et al [2009]. Given that we are observing 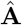 containing observed estimates 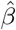, Results 1 and 2 allow for the mean and variance of the perturbation 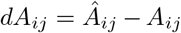 to be defined, with full proof given in the Appendix.

#### Result 1

Given **Q**, if 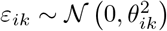 and 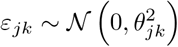 are independent ∀*i, j, k*, then

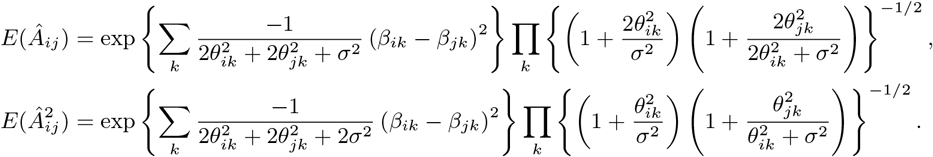

#### Result 2

Given **Q**, if 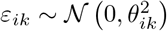 and 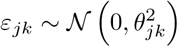 are independent ∀*i, j, k*, then for large values of the dimension *n*,

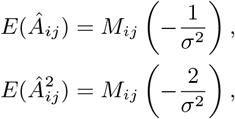

where 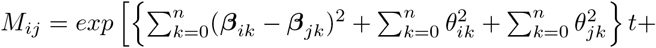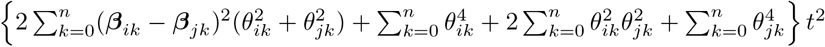, the moment generating function of the normal distribution with given mean and variance.

We next consider the perturbation on the Laplacian matrix. In our setting, we observed 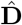 rather than **D**, thus we have 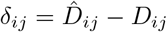. Therefore we observe the perturbation on the Laplacian matrix defined by 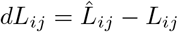.

#### Result 3

Given that perturbations *d***A** and *d***L** are small compared to **A** and **L**, respectively, then,

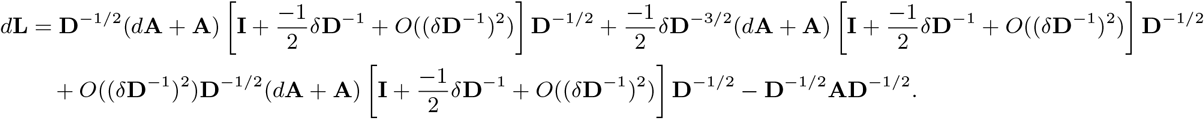

One can directly estimate the mean and variance of *d***L** from this form.

### 4.3 Adjustment for error

Due to the estimation of the regression coefficients used in regression-based spectral clustering, we obtain biased estimates of the elements of the affinity matrix. We proceed to adjust the algorithm to account for this observed affinity matrix, by considering the expectation of the estimate to effectively unbias the affinity matrix. Recall that we obtain the regression coefficient estimates 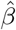 with associated error 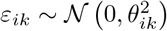, where we observe the estimated variance 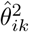. We will seek to evaluate the expectation of the affinity matrix in order to be able to calculate an adjusted, unbiased estimate of its elements without making an equal variance assumption across elements. Therefore given the form of the elements of the affinity matrix **A** as defined in Section 4.2, we consider the relationship,

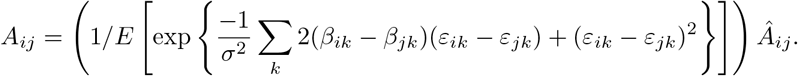

Towards defining the expectation, recall that 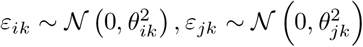 are independent. Then,

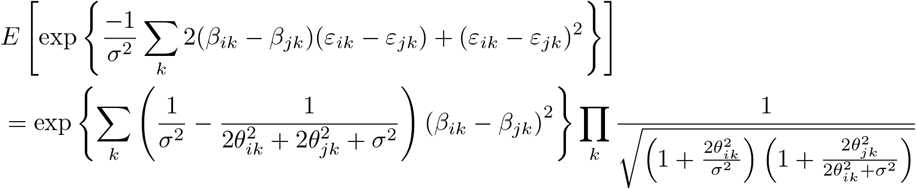

Further details of this expectation are given in the Appendix, where we leverage the independence of *ε*_*ik*_ and *ε*_*ij*_ to obtain the closed form. Therefore we have an adjusted estimate in the affinity matrix given by,

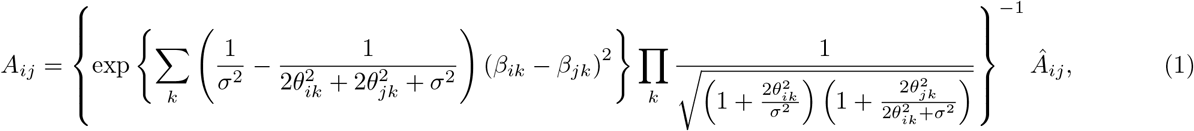

from which we can use estimates from the regression model and proceed with the standard steps of spectral clustering.

### 4.4 Selection of *σ*^2^

The spectral clustering algorithm of Ng, Jordan, and Weiss [2002] requires the scaling parameter *σ*^2^, as noted in Section 4.1. The scaling parameter is a measure of the similarity between nodes and impacts the calculation of the affinity matrix. The authors propose a search over a manually specified space for the value of *σ*^2^ that leads to the tightest clusters after the final k-means clustering step [13]. This requires one to repeatedly run the spectral clustering algorithm to assess the performance of *σ*^2^ values while simultaneously considering the number of clusters *k* in the data.

In order to reduce the computational burden and lack of data-driven automation in this selection procedure, Zelnik-Manor and Perona [2005] proposed a highly specific scaling parameter based the neighborhood of a given node *s*_*i*_ [29]. They propose to compute a local scale parameter *σ*_*i*_ for each data point as *σ*_*i*_ = *d*(*s*_*i*_, *s*_*K*_), where *d*(·) is a distance measure and *s*_*K*_ is the *K* nearest neighbor of *s*_*i*_ with *K* standardly taken to be equal to 7. They then redefine *σ*^2^ for each *A*_*ij*_ in the spectral clustering algorithm as *σ*_*i*_*σ*_*j*_.

We seek to take an principled approach to selecting *σ*^2^ while maintaining computational efficiency. The Zelnik-Manor and Perona approach is demonstrated to perform well when the data is on different scales; however, we note that in our setting of an eQTL network the features are all of the same data type and on the same scale. We thus use the first step proposed in Zelnik-Manor and Perona [2005] to compute the distance metric for each data point and its neighbors. We then reduce the computational burden by taking the median of the distances calculated for each paired point and *K* nearest neighbor, as given below. In practice, we observed that for our data setting the distance metric at the nearest neighbor does not vary highly across *i*; however, this performance would decrease in a setting where we consider a network of features with different scaling.

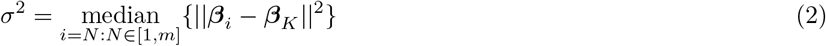

## Acknowledgements

This work was supported by the National Science Foundation Graduate Research Fellowship (DGE1144152) and by the National Institutes of Health F31 (HL138832-01). This work was conducted under dbGaP approved protocol 9112.

## References

[1] Girvan, M. and Newman, M. E. (2002). Community structure in social and biological networks. Proceedings of the national academy of sciences 99, 7821–7826.

[2] Sonawane, A. R., Weiss, S. T., Glass, K., and Sharma, A. (2019). Network medicine in the age of biomedical big data. arXiv preprint arXiv:1903.05449.

[3] Fagny, M., Paulson, J. N., Kuijjer, M. L., Sonawane, A. R., Chen, C.-Y., Lopes-Ramos, C. M., Glass, K., Quackenbush, J., and Platig, J. (2017). Exploring regulation in tissues with eqtl networks. Proceedings of the National Academy of Sciences pp. 201707375.

[4] Platig, J., Castaldi, P. J., DeMeo, D., and Quackenbush, J. (2016). Bipartite community structure of eqtls. PLoS computational biology 12, e1005033.

[5] Fortunato, S. (2010). Community detection in graphs. Physics reports 486, 75–174.

[6] Newman, M. E. (2006). Modularity and community structure in networks. Proceedings of the national academy of sciences 103, 8577–8582.

[7] Sneath, P. H., Sokal, R. R., et al. (1973). Numerical taxonomy. The principles and practice of numerical classification.).

[8] Tamayo, P., Slonim, D., Mesirov, J., Zhu, Q., Kitareewan, S., Dmitrovsky, E., Lander, E. S., and Golub, T. R. (1999). Interpreting patterns of gene expression with self-organizing maps: methods and application to hematopoietic differentiation. Proceedings of the National Academy of Sciences 96, 2907–2912.

[9] Friedman, J., Hastie, T., and Tibshirani, R. (2001). The elements of statistical learning volume 1. (Springer series in statistics New York).

[10] Weiss, Y. (1999). Segmentation using eigenvectors: a unifying view. In The proceedings of the seventh IEEE international conference on Computer vision volume 2 IEEE pp. 975–982.

[11] Rohe, K., Chatterjee, S., and Yu, B. (2011). Spectral clustering and the high-dimensional stochastic blockmodel. The Annals of Statistics pp. 1878–1915.

[12] Newman, M. E. (2013). Spectral methods for community detection and graph partitioning. Physical Review E 88, 042822.

[13] Ng, A. Y., Jordan, M. I., and Weiss, Y. (2002). On spectral clustering: Analysis and an algorithm. In Advances in neural information processing systems pp. 849–856.

[14] Shi, J. and Malik, J. (2000). Normalized cuts and image segmentation. IEEE Transactions on Pattern Analysis and Machine Intelligence 22, 888–905.

[15] Meila, M. and Shi, J. (2001). A random walks view of spectral segmentation. In 8th International Workshop on Artificial Intelligence and Statistics (AISTATS).

[16] Speer, N., Frohlich, H., Spieth, C., and Zell, A. (2005). Functional grouping of genes using spectral clustering and gene ontology. In Proceedings. 2005 IEEE International Joint Conference on Neural Networks, 2005. volume 1 IEEE pp. 298–303.

[17] Wang, B., Mezlini, A. M., Demir, F., Fiume, M., Tu, Z., Brudno, M., Haibe-Kains, B., and Goldenberg, A. (2014). Similarity network fusion for aggregating data types on a genomic scale. Nature methods 11, 333.

[18] Wang, Y. R. and Huang, H. (2014). Review on statistical methods for gene network reconstruction using expression data. Journal of theoretical biology 362, 53–61.

[19] Kluger, Y., Basri, R., Chang, J. T., and Gerstein, M. (2003). Spectral biclustering of microarray data: coclustering genes and conditions. Genome research 13, 703–716.

[20] Inoue, K., Li, W., and Kurata, H. (2010). Diffusion model based spectral clustering for protein-protein interaction networks. PloS one 5, e12623.

[21] Von Luxburg, U. (2007). A tutorial on spectral clustering. Statistics and computing 17, 395–416.

[22] Barabási, A.-L. et al. (2016). Network science. (Cambridge university press).

[23] Carrington, P. J., Scott, J., and Wasserman, S. (2005). Models and methods in social network analysis volume 28. (Cambridge university press).

[24] Huang, L., Yan, D., Taft, N., and Jordan, M. I. (2009). Spectral clustering with perturbed data. In Advances in Neural Information Processing Systems pp. 705–712.

[25] Consortium, G. et al. (2015). The genotype-tissue expression (gtex) pilot analysis: Multitissue gene regulation in humans. Science 348, 648–660.

[26] Paulson, J. N., Chen, C.-Y., Lopes-Ramos, C. M., Kuijjer, M. L., Platig, J., Sonawane, A. R., Fagny, M., Glass, K., and Quackenbush, J. (2017). Tissue-aware rna-seq processing and normalization for heterogeneous and sparse data. BMC bioinformatics 18, 437.

[27] Hicks, S. C., Okrah, K., Paulson, J. N., Quackenbush, J., Irizarry, R. A., and Bravo, H. C. (2017). Smooth quantile normalization. Biostatistics 19, 185–198.

[28] Hartigan, J. A. and Wong, M. A. (1979). Algorithm as 136: A k-means clustering algorithm. Journal of the Royal Statistical Society. Series C (Applied Statistics) 28, 100–108.

[29] Zelnik-Manor, L. and Perona, P. (2005). Self-tuning spectral clustering. In Advances in neural information processing systems pp. 1601–1608.

